# A dynamic subpopulation of CRISPR-Cas overexpressers allows *Streptococcus pyogenes* to rapidly respond to phage

**DOI:** 10.1101/2024.01.11.575229

**Authors:** Marie J. Stoltzfus, Rachael E. Workman, Nicholas C. Keith, Joshua W. Modell

## Abstract

Many CRISPR-Cas systems, which provide bacteria with adaptive immunity against phages, are transcriptionally repressed in their native hosts. How CRISPR-Cas expression is induced as needed, for example during a bacteriophage infection, remains poorly understood. In *Streptococcus pyogenes*, a non-canonical guide RNA *tracr-L* directs Cas9 to autorepress its own promoter. Here, we describe a dynamic subpopulation of cells harboring single mutations that disrupt Cas9 binding and cause CRISPR-Cas overexpression. Cas9 actively expands this population by elevating mutation rates at the *tracr-L* target site. Overexpressers exhibit higher rates of memory formation, stronger potency of old memories, and a larger memory storage capacity relative to wild-type cells, which are surprisingly vulnerable to phage infection. However, in the absence of phage, CRISPR-Cas overexpression reduces fitness. We propose that CRISPR-Cas overexpressers are critical players in phage defense, enabling bacterial populations to mount rapid transcriptional responses to phage without requiring transient changes in any one cell.

## INTRODUCTION

Effective immune systems must quickly identify and destroy foreign threats while avoiding similar motifs within their host. Bacteria encode a growing set of immune effectors that defend against bacteriophages (phages) and plasmids, but how these systems balance immunity and autoimmunity is still an open question. CRISPR-Cas systems, which provide bacteria with adaptive immunity against foreign nucleic acids, have been coopted as transformative gene-editing tools, but in many cell types, heterologous overexpression of Cas nucleases is toxic^1–4^. In their native hosts, CRISPR-Cas systems are often transcriptionally repressed in the absence of phage or other stressors. While this repression could mitigate autoimmunity, it remains unclear (i) whether native CRISPR-Cas promoters are strong enough to cause autoimmunity in their de-repressed state and (ii) how CRISPR-Cas expression can be transiently induced as needed.

In some bacterial and archaeal species, CRISPR-Cas expression increases in direct response to a phage infection^5–9^. However, any response to a phage infection is a race against the relatively short lytic cycle, which may limit the utility of such a response. An alternative strategy is to increase CRISPR-Cas expression *before* the phage arrives. Indeed, many CRISPR-Cas repressors are regulated by environmental signals that may portend phage infection, including population density, envelope stress, and nutrient availability^10–13^. However, phage infection may or may not be preceded by these signals, and we wondered whether there could be a more reliable mechanism to prepare cells for a phage infection.

CRISPR-Cas immunity comprises three stages: adaptation, biogenesis, and interference. During adaptation in the *Streptococcus pyogenes* type II-A system, a 30 bp piece of phage DNA, or “spacer,” is captured from a phage and incorporated into the 5’ end of the CRISPR array between flanking 36 bp repeat sequences^14–16^. During biogenesis, the molecular memories within the CRISPR array are transcribed into a single precursor CRISPR RNA (*pre-crRNA*) and a transactivating *crRNA* (*tracrRNA*) facilitates processing of the *pre-crRNA* into short CRISPR RNAs (*crRNA*s), each carrying the sequence from one spacer^17–19^. Finally, during interference, the *tracrRNA*:*crRNA* duplex directs the nuclease Cas9 to cleave dsDNA complementary to the *crRNA* spacer^19–21^. The *crRNA* target must be next to a 5’-NGG-3’ protospacer adjacent motif (PAM) to activate Cas9 catalytic activity^22–24^. The CRISPR array, where spacer sequences are stored within the genome, lacks PAM sequences, thus preventing targeting of self-DNA.

In *S. pyogenes,* CRISPR-Cas expression is regulated by a long-form *tracrRNA* (*tracr-L*)^25^. *tracr-L* folds into a single-guide RNA that repurposes Cas9 into a transcriptional repressor of the *cas* operon promoter (P*_cas_*), which encodes Cas9 and three additional Cas proteins which are required only for adaptation (Fig. 1A). In a heterologous *Staphylococcus aureus* host, the loss of *tracr-L* results in enhanced CRISPR-Cas expression and phage defense, but CRISPR-Cas overexpressers also suffer a fitness cost in the absence of phage. We hypothesized that dynamic regulation of CRISPR-Cas expression could allow cells to defend against phage while mitigating autoimmunity.

**Figure 1.**
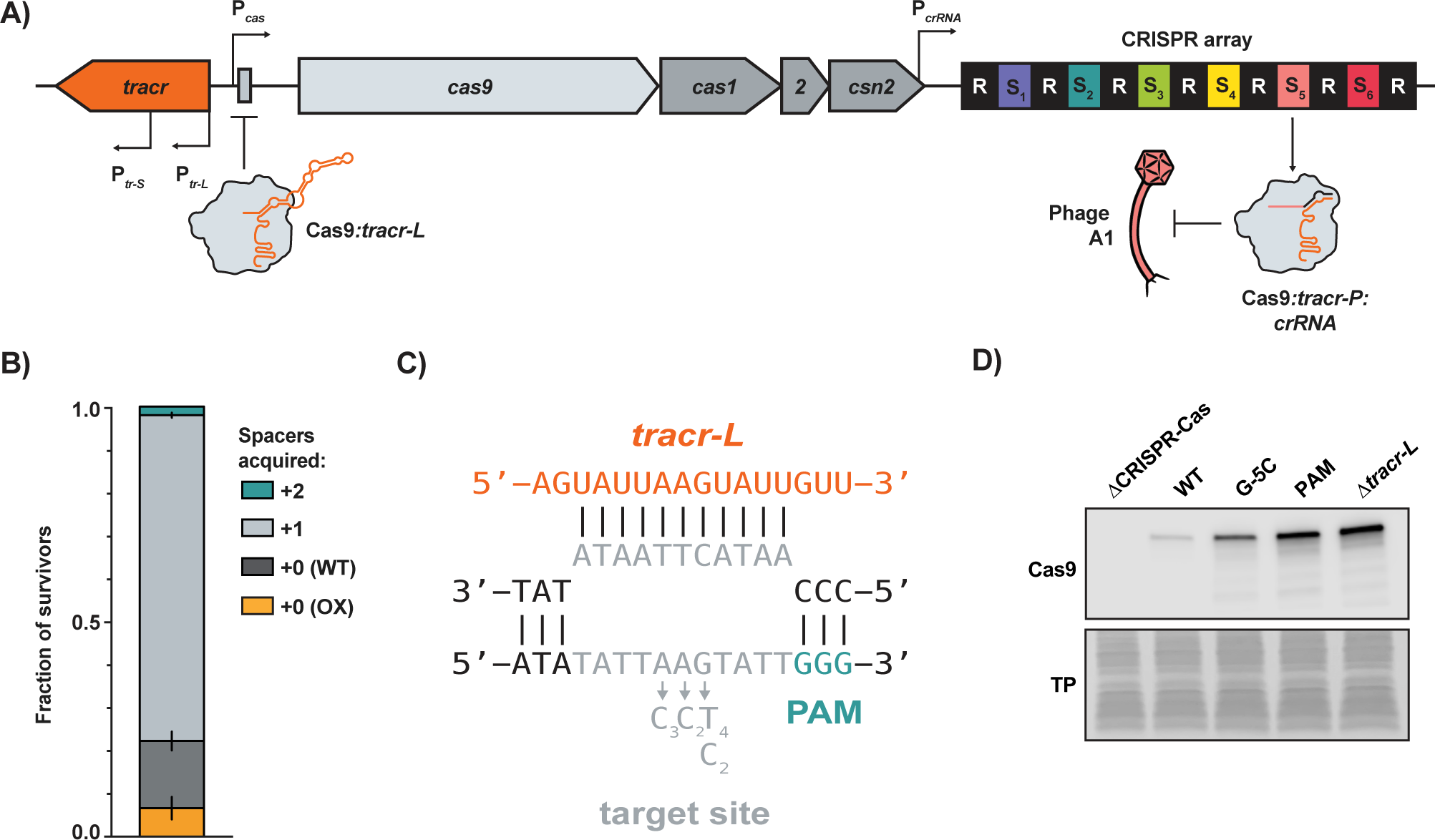
A subpopulation of CRISPR-Cas overexpressers expands during phage infection. **A)** The type II-A *S. pyogenes* CRISPR-Cas system. The repressive Cas9:*tracr-L* complex binds downstream from the *cas* operon transcriptional start site. Spacer 5 targets phage A1. **B)** *S. pyogenes* cells were infected with phage A1 at MOI = 1 in soft agar and the genotype of surviving colonies was determined by PCR and Sanger Sequencing (OX, overexpresser with mutation in the P*_cas_* target site). Error bars are standard error. n=3. **C)** The R-loop formed between *tracr-L* (orange) and the P*_cas_* target site (grey). Mutations in the target site observed in survivors of the infection described in (B) are shown. Subscripts indicate the number of individual survivors with each mutation. **D)** Representative Western blot probing for Cas9 levels in stationary phase cultures of *S. pyogenes* with the indicated mutations in P*_cas_* (G-5C, PAM) or a *tracr-L* deletion (TP, Ponceau stain for total protein).

Here, we show that a subpopulation of cells harbors a single mutation in P*_cas_* that disrupts Cas9:*tracr-L* repression and leads to CRISPR-Cas overexpression *before* the arrival of phage. Overexpressers exhibit higher rates of interference with spacers across the CRISPR array and higher rates of immunization when challenged with a novel phage. Additionally, unlike WT cells which can efficiently interfere with only a small number of spacers, overexpressers can maintain a larger repertoire of potent spacers. For these reasons, the overexpressing subpopulation is enriched following phage infection. However, in the absence of phage, the overexpressing subpopulation shrinks due to fitness costs. Finally, we show that Cas9:*tracr-L* binding results in higher rates of mutation within the *tracr-L* binding site and PAM, thus expanding the size of the overexpressing population. Altogether, we propose a model whereby the maintenance of a mutant subpopulation of CRISPR-Cas overexpressers enables the greater bacterial population to rapidly respond to phage without requiring a transient response in any single cell.

## RESOURCE AVAILABILITY

### Data Availability

Data is available upon request from corresponding author, Joshua Modell.

### Code Availability

Code is available upon request from corresponding author, Joshua Modell.

### Material Availability

All materials are available upon request without restriction from corresponding author, Joshua Modell.

## METHODS

### Strains

All experiments were performed in the *Streptococcus pyogenes* M1 GAS strain SF370^26^ or *Staphylococcus aureus* RN4220 and derivatives. Plasmid cloning was performed in *Escherichia coli* DH5α or RN4220.

### Culture Conditions

*S. pyogenes* and *S. aureus* strains were maintained in Bacto Brain Heart Infusion (BHI) broth unless otherwise stated, and *E. coli* strains in LB broth. Liquid cultures of *S. pyogenes* strains were grown at 37°C without shaking, and liquid cultures of *S. aureus* and *E. coli* were grown shaking at 220 rpm at 37°C. Antibiotics for strain construction and plasmid maintenance were supplemented at the following concentrations for *S. pyogenes*: chloramphenicol (cm), 3 µg/mL; spectinomycin (spec), 100 µg/mL. For *S. aureus*: cm, 10 µg/mL; spec, 250 µg/mL; rifampicin (rif), 0.5 µg/ml. For *E. coli*: cm, 25 µg/mL; spec, 50 µg/mL; kanamycin (kan), 50 µg/mL.

For phage assays, overnight cultures of *S. pyogenes* cells were grown in THY (Todd Hewitt Broth (BD 249240) + 2% Bacto Yeast Extract (Gibco 212750)), and outgrowths from stationary phase in THY dialysate (THY-D, see below) supplemented with 5 mM CaCl_2_ and 2 mg/mL NaHCO_3_.

### Todd-Hewitt Dialysate Media (THY-D)

To prepare media for *S. pyogenes* phage infections as described previously^27^, 30 g Bacto Todd Hewitt Broth media and 20 g Bacto yeast extract in 100 ml dH_2_O were microwaved and agitated until homogenous, then poured into 11 inches of dialysis tubing (Spectra Por S/P 3; molecular weight cut-off 3.5 kDa) and sealed. The dialysis bag was floated in 2 L dH_2_O for 1 hour at room temperature (RT), then the water was replaced and the media dialyzed for 2 more hours. The bag was then transferred to a beaker with 4 L fresh dH_2_O and dialyzed at 4°C for 16 hrs. Media was transferred to a bottle, brought to 1 L final volume, and autoclaved for 45 min.

### Phages

Phage A1 and derivatives were propagated in C13, a derivative of the prophage-cured SF370 strain CEM1Δɸ^28^ that lacks spacers 1-5 (a generous gift from Andrew Varble, University of Rochester), and stored at 4°C in THY-D.

### Plasmid Construction

See supplemental table 3 for oligos and plasmids used in this study. PCR to generate fragments for Gibson assemblies was performed using Phusion HF DNA polymerase and 5X Phusion Green Reaction Buffer (Thermo) per manufacturer’s instructions. Gibson assemblies were performed as described previously^29^. Briefly, equimolar amounts of each PCR product were mixed and brought to 5 µl with dH_2_O, then mixed with 15 µl Gibson master mix. The reaction was incubated at 50°C for 1 hr. Before transformation into *S. aureus*, the reactions were drop dialyzed for 30-60 min, and 5 µl was electroporated into 50 µl competent cells. For transformations into *E. coli*, 5 µl of the Gibson reaction was transformed via heat shock into DH5α. Plasmids were purified using the Spin MiniPrep kit (Qiagen 27104) per manufacturer’s instruction with the following modification for *S. aureus*: lysostaphin (Ambi Products LLC, LSPN-50, 100 µg/mL final concentration) was added to pellets resuspended in P1 and incubated at 37°C for 20 min before the addition of P2.

### Colony Lysis

*S. pyogenes* colonies were resuspended in 1X PBS with PlyC (1 µg/ml final concentration; a generous gift from Chad Euler, Hunter College) and incubated at RT for 10 min followed by 10 min at 98°C. For PCRs, 0.5-2 µl of lysate was used as template.

### Genomic DNA Extractions

DNA was extracted from *S. pyogenes* using the DNeasy Blood and Tissue Kit (Qiagen 69504) with the following modifications: cell pellets were resuspended in 180 µl PBS supplemented with 1 µg/mL PlyC and incubated at RT for 20 min before proceeding with the kit protocol beginning at proteinase K digestion.

### Transformations into *S. pyogenes*

To prepare electrocompetent cells, *S. pyogenes* at OD_600_ 0.4 were pelleted at 4°C for 20 min at 3200 rcf. The pellet was washed with one volume ice-cold 10% glycerol, then with 1/20^th^ volume, and resuspended in 1/150^th^ volume 10% glycerol. Cells were used immediately or stored at −80°C.

Plasmids were drop dialyzed in dI water for 45 min and 0.3–1 µg were mixed with 50 µl competent cells on ice. The cell/plasmid mixture was transferred to a chilled Gene Pulser 0.1 cm Cuvette (Bio-Rad 165-2089) and electroporated in a Gene Pulser Xcell (Bio-Rad; 2.5 kV/cm, 200 Ω, 25 µF). 950 µl BHI was added, and the transformed cells were incubated at 37°C for 1-2 hrs before plating on appropriate antibiotic selection plates and incubating at 37°C for 18-24 hrs.

### Allelic Exchange in *S. pyogenes*

To generate mutations in the *S. pyogenes* chromosome, 500-1,000 bp homology arms surrounding the desired mutation were inserted into the temperature-sensitive pCRS backbone via Gibson assembly, then transformed into and propagated in *E. coli* DH5a. Oligo sequences and plasmids are listed in supplemental table 3. Allelic exchange constructs were transformed into *S. pyogenes* as described above, except that outgrowths and incubations were performed at 30°C to permit plasmid replication and plates were incubated for 36-48 hrs to allow colonies to form. Single colonies were inoculated into prewarmed BHI + spec and incubated overnight at 37°C, then plated on antibiotic at 37°C and incubated overnight to select for single integrants. Single colonies were restruck on antibiotic at 30°C and incubated overnight, then inoculated into BHI without antibiotic and incubated overnight at 30°C to allow for plasmid excision and loss. Cultures were diluted 1:1,000 in prewarmed BHI and incubated at 37°C for 8-12 hrs twice, then plated on BHI agar and incubated overnight at 37°C. To screen for double crossovers, 80-120 colonies were patched onto plates with and without antibiotic, and antibiotic-sensitive colonies were further screened for the desired mutation via PCR and Sanger sequencing.

### Phage Engineering

To generate mutations in phage A1, a plasmid containing the desired mutation with ∼500 bp flanking homology regions was transformed into C13. Phage A1 was propagated on this strain to allow for recombination, and recombinant phages were selected by plating to single plaques on C13 Δ*tracr-L* harboring a plasmid with a spacer targeting WT, but not mutant, phage. Individual plaques were screened for the correct mutation via PCR and Sanger sequencing.

### Phage Soft Agar Assays

200 µl *S. pyogenes* at OD_600_ 0.3-0.6 in THY-D + 5 mM CaCl_2_ and 2 mg/mL NaHCO_3_ and phage were mixed with 2 mL THY-D + 0.2% agarose supplemented with 5 mM CaCl_2_ and 2 mg/mL NaHCO_3_ and plated on BHI 1.5% agar plates. Plates were allowed to set at RT for 20 min before incubating overnight at 37°C. To save phage survivors as new strains for further experiments, colonies from soft agar plates were first restruck to single colonies twice to dilute residual phage.

### Phage Plaque Forming Unit Assays

200 µl *S. pyogenes* at OD_600_ 0.6 in THY-D + 5 mM CaCl_2_ and 2 mg/mL NaHCO_3_ were mixed with 2 mL THY-D + 0.2% agarose + 5 mM CaCl_2_ and 2 mg/mL NaHCO_3_ and plated on BHI 1.5% agar plates. After plates set for 20 min at RT, 2-3 µl of 10-fold dilutions of phage in THY-D were spotted onto the plates. In some cases, plates were tilted gently, allowing phage to drip down the plate to separate plaques for more accurate quantification. After 20 min at RT, plates were incubated overnight at 37°C.

### Phase Microscopy for MOI Correction

*S. pyogenes* forms long chains in liquid culture. Therefore, traditional colony forming unit (CFU) enumeration techniques can underestimate the number of viable cells in a culture due to a chain, which contains multiple cells, forming only one colony. To determine the average chain length in *S. pyogenes* cultures, cells were outgrown to OD_600_ 0.5 in Thy-D with 5 mM CaCl_2_ and 2 mg/mL NaHCO_3_. Cultures were diluted 10-fold and spotted onto agarose pads (1% UltraPure™ Agarose (Invitrogen) in 1X PBS). Cells were imaged on an IX83 inverted microscope with a 100X phase oil immersion objective (Olympus), and 7-10 fields of view were captured for each of three biological replicates. The average chain length under these growth conditions was determined to be 10 cells/chain. This correction factor was included in MOI calculations as follows (PFU, plaque forming units):

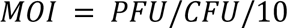

### Western Blots

*S. pyogenes* cell pellets were resuspended in 1X PBS supplemented with 1 µg/mL PlyC and incubated at RT for 10 min. One volume 2X Laemmli solution (Bio-rad, 1610737) supplemented with 1.3 mM ß-mercaptoethanol (ThermoFisher, 21985023) was added and incubated at 98°C for 10 min. Samples were centrifuged at 6,000 rcf for 1 min, and the cleared lysate was loaded onto 4-20% Tris-glycine gels (MINI-Protean TGX Pre-cast, Bio-Rad, 4561095) and electrophoresed at 200 V for 30 min. Protein was transferred to a PVDF membrane (hydrated in methanol and prewet in 1X Transfer Buffer) between blotting paper stacks in the Trans-blot Turbo (Bio-Rad; 1.6 A for 10 min). The membrane was cut at the 70 kDa ladder marker, and the lower molecular weight piece was stained with Ponceau (Sigma, P3504) for 1 hour then washed with dI water for total protein visualization. The higher molecular weight piece of the membrane was blocked in 5% nonfat dry milk in TBST for 1 hr, then probed with a 1:1,000 dilution of Cas9 monoclonal antibody (Cell Signaling, 7A9-3A3) overnight at 4°C. After washing three times for 10 min each with TBST buffer at RT, the membrane was incubated in a 1:10,000 dilution of Goat anti-mouse HRP-conjugated secondary antibody (Pierce, PA174421) for 1 hr at RT. The membrane was washed three more times, then visualized on the Odyssey Fc (LICOR) using Clarity Western ECL Substrate (Bio-Rad).

### Plasmid Targeting Assay

Electrocompetent *S. pyogenes* cells (50 µl) were mixed with 300 ng plasmid and electroporated as described above. After 1 hr at 37°C, 100 µl of an appropriate dilution to produce single colonies was plated on plain and cm plates using sterile glass beads. Plates were incubated for 18 hrs (plain) or 36 hrs (cm) before counting CFUs. Transformation efficiencies were calculated for each plasmid/strain combination and normalized to the transformation efficiency of an EV control.

### Mutagenesis Assay

To determine if Cas9:*tracr-L* binding to the bacterial genome is mutagenic, a strain of *S. aureus* RN4220 was constructed with the SF370 CRISPR-Cas system integrated into a genomic safe harbor^30^. The CRISPR-Cas system contained the following mutations: Δ*tracr-L*, C-terminal Cas9-GFPmut2 fusion, dCas1, and a single spacer in the CRISPR array targeting the *S. aureus* phage ɸNM4ɣ4. Two plasmids, one on a pC194 backbone and one on pE194, were added containing an IPTG inducible *tracr-L*\*, which is an allele that only produces *tracr-L* due to a mutation in the *tracr-S* promoter. Cells were struck out with or without the addition of IPTG to induce *tracr-L.* Single colonies were inoculated (6 replicates per condition) with the same IPTG treatment as on the plate. Cultures were grown overnight then diluted 1:100 in the morning and again in the evening. The next morning, all cultures were diluted 1:100 in 1 mM IPTG and grown for 5.5 hours to induce *tracr-L* and normalize Cas9 levels between the two conditions. Each culture was then plated in three different ways. To i) determine total CFUs, 10-fold dilutions of each culture were plated on BHI agar supplemented with cm and erm. CFUs were enumerated after 15 hrs at 37°C. To ii) determine relative global mutation rates, 100 µl was plated on rifampicin. CFUs were counted after 15 hrs at 37°C and normalized to total CFUs. To iii) select for overexpressing mutants, 200 ODul of each culture with ɸNM4ɣ4 (MOI 100) were plated in 6 ml soft agar (BHI + 0.75% agar) supplemented with cm, erm, 5 mM CaCl2, and 1 mM IPTG. After setting at RT for 15 min, plates were incubated at 37°C for 36 hrs.

To determine the fraction of colonies that survived phage infection due to mutations in P*_cas_*, ten colonies from each replicate were lysed in colony lysis buffer (250 mM KCl, 5 mM MgCl_2_, 50 mM Tris-HCl pH 9.0, 0.5% Triton X-100) with lysostaphin (Ambi Products LLC, LSPN-50, 60 µg/ml final concentration), followed by PCR amplification and Sanger Sequencing. This fraction was used to determine the fraction of overexpressing phage survivors relative to total CFUs.

### Competitions

Single colonies were inoculated into BHI and incubated overnight at 37°C. Co-cultures of two strains were prepared the next morning by mixing 600 µl of each strain. 1 mL was pelleted, flash frozen in liquid nitrogen, and saved for gDNA extraction (“Day 0”). 100 µl was used to inoculate 10 mL BHI. Cultures were incubated at 37°C for 8 hrs, then diluted 1:1,000 and incubated overnight. Cultures were passaged in the same way for four days (1:100 dilution in the morning, followed by 1:1,000 dilution in the evening), and cell pellets were saved for gDNA extraction on the morning of the second and fourth day. DNA from all saved cell pellets was extracted as described above.

To quantify competitive indices in competitions with Δ*tracr-L* cells, a semi-quantitative PCR was performed. 1 ng of DNA from each time point was amplified with primers spanning the *tracr-L* deletion, and PCR products were run on a 2% agarose gel with 0.5 µg/ml ethidium bromide (90 V, 40 min). Bands were imaged on a Molecular Imager® Gel Doc XR (Bio-Rad). The integrated density of the bands corresponding to WT and Δ*tracr-L* were quantified with ImageJ and, after background subtraction, used to quantify the fraction of Δ*tracr-L* cells remaining.

To quantify competitive indices in competitions with PAM or G-5C overexpressers, a PCR across the *tracr-L* binding site was performed using indexed primers with partial Illumina adaptors. The PCR products were purified with the QIAquick PCR Purification Kit (Qiagen), pooled, and sequenced with Azenta’s Amplicon-EZ service. The resulting reads were demultiplexed and fractions of PAM and G-5C were quantified with custom Python scripts.

### Cas9 ChIP-Seq

Cultures of *S. pyogenes* were grown to mid-logarithmic or stationary phase in BHI at 37°C and diluted to an OD_600_ of 0.4 in 40 ml BHI. Cells were fixed immediately with formaldehyde (1% final concentration) and incubated at RT for 20 min, then quenched with glycine (0.5 M final concentration) and incubated at RT for 10 min. Cells were washed twice with one volume of ice-cold 1X TBS, then lysed by resuspending in FA lysis buffer (50 mM HEPES-KOH pH 7.5, 0.1% sodium deoxycholate, 0.1% SDS, 1mM EDTA, 1% Triton X-100) with 1X cOmplete protease inhibitor cocktail and 1 ug/ml PlyC and incubating at 37°C for 20 min at 300 rpm. Samples were sonicated on ice with a Branson Sonifier 450 (constant duty cycle, output 3, 6 pulses of 15 s) to produce fragments of 250-500 bp. Cell debris was pelleted by centrifugation and the supernatant (lysate) was retained. Samples of each lysate were saved for input control libraries.

To immunoprecipitate Cas9-bound DNA, for each sample, 25 µl Dynabeads Protein G (ThermoFisher) were washed three times in PBS with BSA (5 mg/ml), then nutated in 75 µl PBS BSA with 2 µl Cas9 antibody (Cell Signaling Technology Cas9 (7A9-3A3) Mouse mAb #14697) for 6 hr at 4°C. Antibody-bound beads were resuspended in 1 ml of lysate and incubated at 4°C on a Roto Genie at setting 3.5 for 14 hrs. Beads were washed serially in 1 ml of the following buffers at 4°C: twice in FA lysis buffer with 150 mM NaCl, once in FA lysis buffer with 500 mM NaCl, once in ChIP wash buffer (10 mM Tris HCl pH 8.0, 250 mM LiCl, 0.5% sodium deoxycholate, 0.5% nonidet-P40, 1mM EDTA), and twice in TE (1mM EDTA). Beads were resuspended in 200 µl ChIP elution buffer (1% SDS, 0.1M NaHCO_3_) and 170 µl ChIP elution buffer was added to 30 µl of saved lysate for input controls. Samples were incubated at 65°C for 1 hr at 600 rpm. Beads were pelleted and the supernatant retained. To reverse crosslinks, NaCl was added to 250 mM final concentration and samples incubated at 65°C overnight. Samples were then incubated with RNase A (ThermoFisher, final concentration 50 ng/µl) at 37°C for 1hr, followed by Proteinase K (Qiagen, final concentration 240 ng/µl) at 55°C for 1 hr. DNA was purified using the QIAquick PCR Purification Kit (Qiagen) per manufacturer’s instructions and eluted in 50 µl TE (0.1 mM EDTA).

Multiplexed libraries of input and ChIP samples were prepared for sequencing using the NEBNext® Ultra™ II DNA Library Prep Kit for Illumina® (E7645S) and NEBNext® Multiplex Oligos for Illumina® (Index Primers Sets 1/2, E7335S/E7500S) per manufacturer’s instructions using 1 ng DNA input and 9 cycles of PCR amplification. Libraries were pooled and sequenced on a MiSeq (paired end, 75 cycles) at the Genetic Resources Core Facility. Cas9 ChIP peaks were called using MACS2 (v2.2.9.1)^31^. See supplemental table 1 for MACS2 output tables.

### RNA-Seq

Cultures of *S. pyogenes* were grown to mid-log or stationary phase in BHI at 37°C in biological triplicate. Cell pellets were flash frozen in liquid nitrogen, and RNA was extracted with the Direct-zol™ RNA Miniprep Kit (Zymo Research) per manufacturer’s instructions with the following modification: to lyse cells, pellets were resuspended in 150 µl PBS with PlyC (1 µg/ml final concentration) and incubated 10 min at RT before adding TRI reagent®. RNA was eluted in 50 µl dH_2_O. 3 ug RNA was treated with Promega RQ1 DNase (M6101) then cleaned up with the Monarch® RNA Cleanup Kit (NEB Biolabs, T2050S) and eluted in 30 µl dH_2_O.

Multiplexed libraries were prepared with NEBNext® rRNA Depletion Kit (Bacteria) (E7850L) and NEBNext® Ultra™ II Directional RNA Library Prep Kit for Illumina® (E7760S) using 450 ng total RNA input. PCR amplification was performed for 11 rounds with NEBNext® Multiplex Oligos for Illumina® (Index Primers Sets 1/2, E7335S/E7500S). Libraries were pooled and sequenced on a NovaSeq6000 (SP chip, paired end, 100 cycles) at the Genetic Resources Core Facility. Transcripts were quantified with Salmon (v1.8.0)^32^ and differential expression analysis was performed with DESeq2 (v1.20)^33^ using a fold-change cutoff of 2 and alpha of 0.05. See supplemental table 2 for DESeq2 output tables.

## RESULTS

### A subpopulation of CRISPR-Cas overexpressers expands during phage infection

We previously found that single base mutations in *tracr-L* or its binding site led to significant upregulation of CRISPR-Cas expression and phage defense in a heterologous *S. aureus* host^25^. We hypothesized that similar, naturally occurring mutations could play a major role in phage defense in the native *S. pyogenes* host. However, this would depend on the mutation rate and the benefits and costs of CRISPR-Cas overexpression. To test whether this mutant subpopulation was present and enriched by phage infection, we infected *S. pyogenes* strain SF370 with phage A1^34^, which is targeted by spacer 5 within the native CRISPR array (Fig. 1A). The protection conferred by this spacer is weak, and most SF370 cells infected with A1 in soft agar do not survive at an MOI of 1. Of the surviving cells that formed colonies, the majority acquired one (70%), or occasionally two (2%) new spacers targeting A1 (Fig. 1B, S1A). By sequencing the *tracr-L* regulatory sequences from survivors that did not acquire additional spacers, we found that 9-50% (2-11% of total survivors) had mutations in the *tracr-L* target site in P*_cas_* (Fig. 1B-C, S1B). To characterize additional overexpressing mutants without having to first screen for spacer acquisition, we repeated the infection in a strain incapable of spacer acquisition (Cas1^E220A^, or dCas1) and observed additional mutations within the target site and PAM as well as a single mutation in *tracr-L* (Fig. S1C). We confirmed that these mutations led to overexpression of Cas9 via Western blot (Fig. 1D). Altogether, these data suggest that a preexisting subpopulation of CRISPR-Cas overexpressing cells becomes enriched during a phage infection.

### Old spacers regain potency in CRISPR-Cas overexpressers

Because the population of overexpressers we observed had not acquired a new spacer after infection by A1, we predicted that these colonies survived due to more efficient interference by spacer 5. Indeed, the overexpressers restricted plaque formation by WT A1, but not by a variant with a PAM mutation in the spacer 5 targeted site (A1esc) (Fig. 2A, S2A-B). Because many CRISPR arrays contain tens or hundreds of spacers, we were surprised by the limited potency of spacer 5 against A1 in WT cells. We next sought to establish the strengths of the other spacers in the SF370 array. We engineered A1esc variants targeted by each of the six spacers in the native CRISPR array and determined their efficiency of plaquing (EOP) on WT and overexpressing cells (Fig. 2B). Curiously, in WT cells, only a subset of spacers (1, 3, and 4) confer significant protection, while the remaining spacers (2, 5, and 6) are relatively weak, even against plasmid targets (Fig. S2C). The limited protection provided by the oldest spacers (5 and 6) is in line with previous work showing that the newest spacers, which reside proximal to the *crRNA* leader, provide more robust interference when the *S. pyogenes* CRISPR system is heterologously expressed on a plasmid in *E. coli* or *S. aureus* cells^16,35^. In the CRISPR-Cas overexpressers, however, all six spacers remain potent, with interference levels approaching the limit of detection of this assay for both a target site mutant (G-5T) and Δ*tracr-L* (Fig. 2B, S2C). Thus, unlike WT cells, CRISPR-Cas overexpressers can utilize the entire native CRISPR array for protection against older (spacers 5 and 6) and poorly targeted (spacer 2) threats.

**Figure 2.**
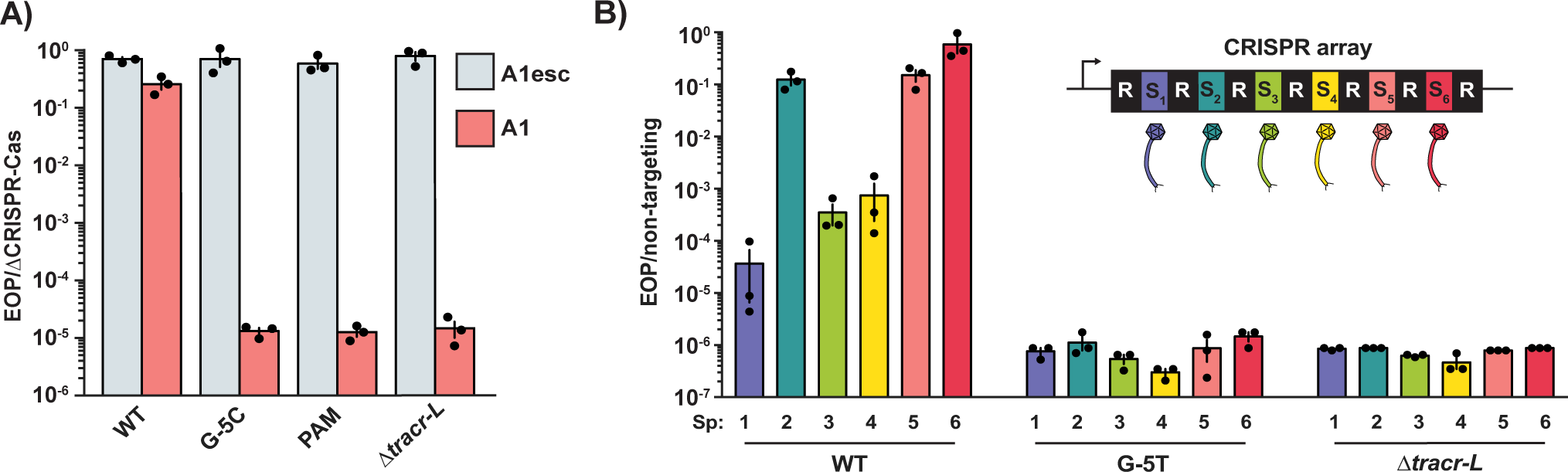
Old spacers regain potency in CRISPR-Cas overexpressers. **A)** Efficiency of plaquing of a targeted phage (A1) and a non-targeted mutant (A1esc) on *S. pyogenes* with the indicated CRISPR-Cas overexpressing mutations. Efficiency of plaquing (EOP) was calculated relative to a ΔCRISPR-Cas strain. **B)** Phages targeted by each of the 6 spacers within the native CRISPR array were plated on WT and CRISPR-Cas overexpressing cells. EOP was calculated relative to a non-targeting strain with all spacers in the CRISPR array deleted. Errors bars are standard error. n=3.

### CRISPR-Cas overexpressers acquire spacers more frequently

We wondered if, in addition to the enhanced potency of existing spacers, CRISPR-Cas overexpressers also acquire new spacers at elevated rates. To test this, we performed an “immunization assay” and infected cells with the non-targeted phage A1esc. To survive infection by this phage, cells must acquire a new spacer, a rare event which occurs in ∼1/1 million infected WT cells under these conditions. In CRISPR-Cas overexpressing cells, immunization rates were ∼60-fold higher than WT (Fig. 3A, S3A). Additionally, overexpressers frequently acquired two or more spacers (17% in G-5C target site mutant; 25% in PAM mutant; 33% in Δ*tracr-L*; Fig. 3B) unlike WT cells (7%). Interestingly, we found that overexpressers occasionally acquired spacers with non-canonical PAMs, including NAG, GGN, NNGG, and NTG (Fig. S3B-C). Although spacers with these PAMs can offer some level of protection^36–38^, Cas9 cleaves these targets less efficiently than those with NGG PAMs. Indeed, spacers with non-canonical PAMs were typically found in arrays with two or more novel spacers, with at least one using a canonical NGG PAM. However, in two cases, an overexpresser acquired a single spacer with a non-canonical NAG or NTG PAM, indicating that they are strong enough to provide protection in a CRISPR-Cas overexpressing background. Spacers with non-canonical PAMs were never observed in WT survivors, and we hypothesize that (i) they may not offer sufficient protection when the CRISPR-Cas system is repressed, or (ii) they may require a second, functional spacer to provide full defense, which occurs less frequently in WT cells.

**Figure 3.**
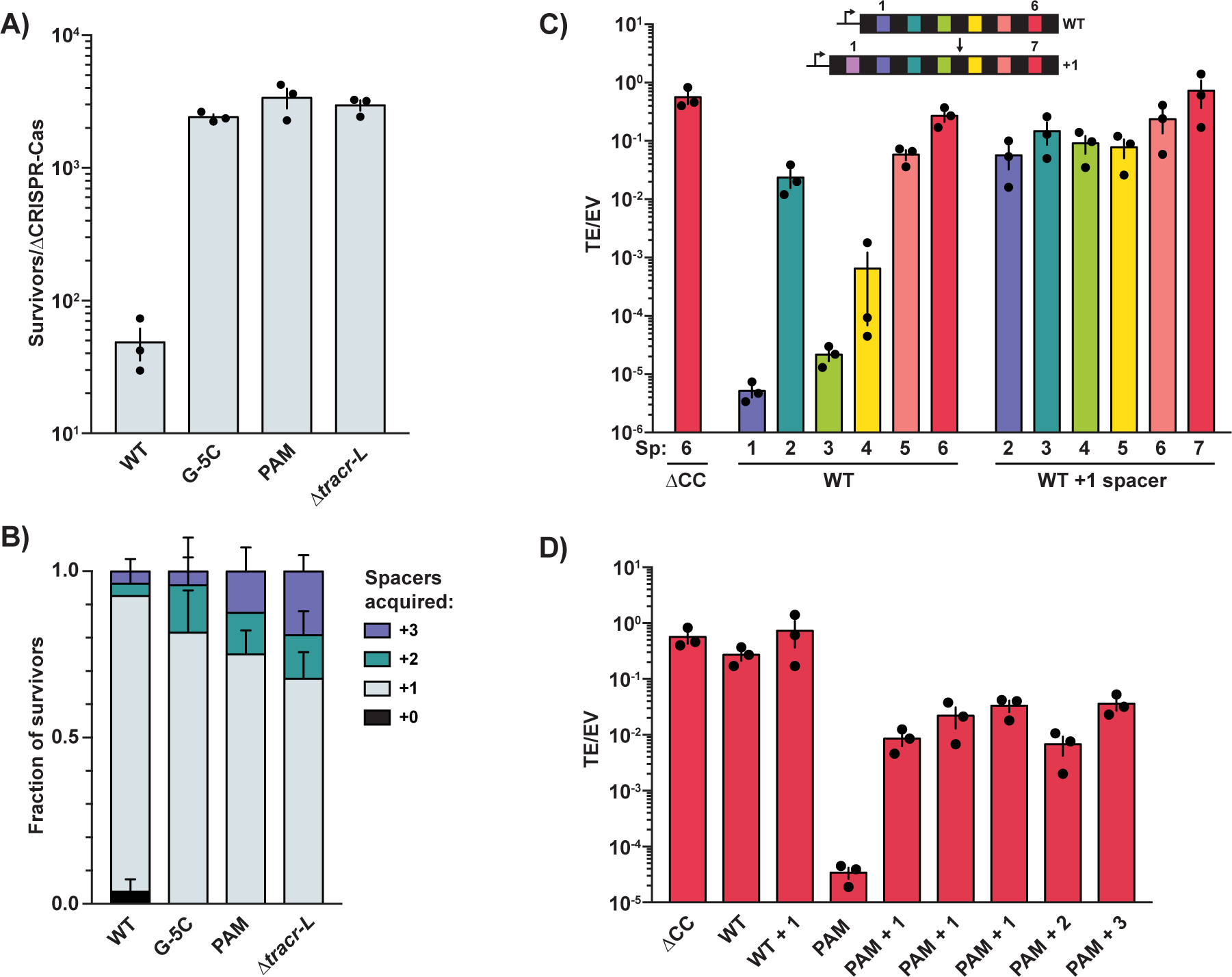
CRISPR-Cas overexpressers generate greater functional spacer diversity. **A)** Survival rates of *S. pyogenes* infected with non-targeted phage A1esc at MOI 5 in soft agar relative to ΔCRISPR-Cas. **B)** Genotypes of survivors from (A) determined by PCR across the CRISPR array. **C)** WT cells and cells that acquired one additional spacer were transformed with plasmids targeted by the indicated spacer (“Sp”) in the WT array. Transformation efficiency was calculated relative to an empty vector control. The leader-proximal spacer is labeled as spacer 1. **D)** PAM mutants that acquired 1-3 additional spacers during infection with A1esc were transformed with a plasmid targeted by spacer 6 in the WT array. Three independent PAM mutants that acquired one spacer were tested. Transformation efficiency is shown, along with the parent strain (PAM). WT and ΔCRISPR-Cas data from (B) are included for comparison. Errors bars are standard error. n=3.

### CRISPR-Cas overexpressers can utilize larger CRISPR arrays

We next asked to what degree CRISPR-Cas overexpression increases the number of functional spacer positions within the CRISPR array. Our prior results suggested that the effective array size in WT cells is limited to the first 4 positions (Fig. 2B), but it was also possible that spacers 5 and 6 are weak due to sequence-specific characteristics. To distinguish between these possibilities, we swapped the first and last spacer in the native array and tested the potency of those spacers in a plasmid targeting assay. We observed that moving the oldest spacer to the leader-proximal position dramatically increased interference efficiency, while moving the newest, most potent spacer to the end of the array severely decreased targeting (Fig. S4D-E). This suggests that the limited utility of old spacers in the native array is due to their position rather than purely sequence-specific differences in cleavage efficiency.

Given that spacers in WT arrays fail to provide robust defense after position 4, we next sought to determine the functional length of arrays in overexpressers. To test this, we performed a plasmid targeting assay in survivors of phage infections that had acquired additional spacers. In WT cells, the acquisition of a single additional spacer led to significantly decreased interference by the spacers that were originally potent in positions one, three, and four (Fig. 3C). In a CRISPR-Cas overexpresser with a mutation in the P*_cas_* target site PAM, even spacer 6 still reduced plaque formation by ∼2 logs after acquisition of three additional spacers, pushing the functional array size to at least position 9 (Fig. 3D). These results highlight the surprising limitations of spacer utility in WT cells and the expanded effective array lengths of CRISPR-Cas overexpressers.

### CRISPR-Cas overexpression comes with a fitness cost

While CRISPR-Cas overexpression provides a clear benefit during phage predation, previous work has shown that high levels of SpyCas9 and other CRISPR-Cas components are toxic in many bacteria^25,39–42^. Because all of these studies involve strong CRISPR-Cas overexpression from multi-copy plasmids, we wondered if CRISPR-Cas overexpression from the genome in naturally occurring *S. pyogenes* mutants is similarly toxic. To test if CRISPR-Cas overexpression decreases fitness in the absence of phage pressure, we performed competition experiments between WT SF370 and cells with mutations in the *tracr-L* target site or PAM sequences (Fig. 4A). We found that overexpressers are indeed less fit than wild type cells (Fig. 4B-C), showing a ∼2 fold reduction over the course of 4 days of passaging. During these experiments, we occasionally observed outliers in which the overexpressing population dominates the culture by day 4 (see Fig. 4B). However, these outliers are rare (see Fig. S4F for additional replicates of the WT vs. PAM competition) and in all cases further examined are accompanied by secondary mutations that lead to elevated growth rates independent of CRISPR-Cas expression (data not shown).

**Figure 4.**
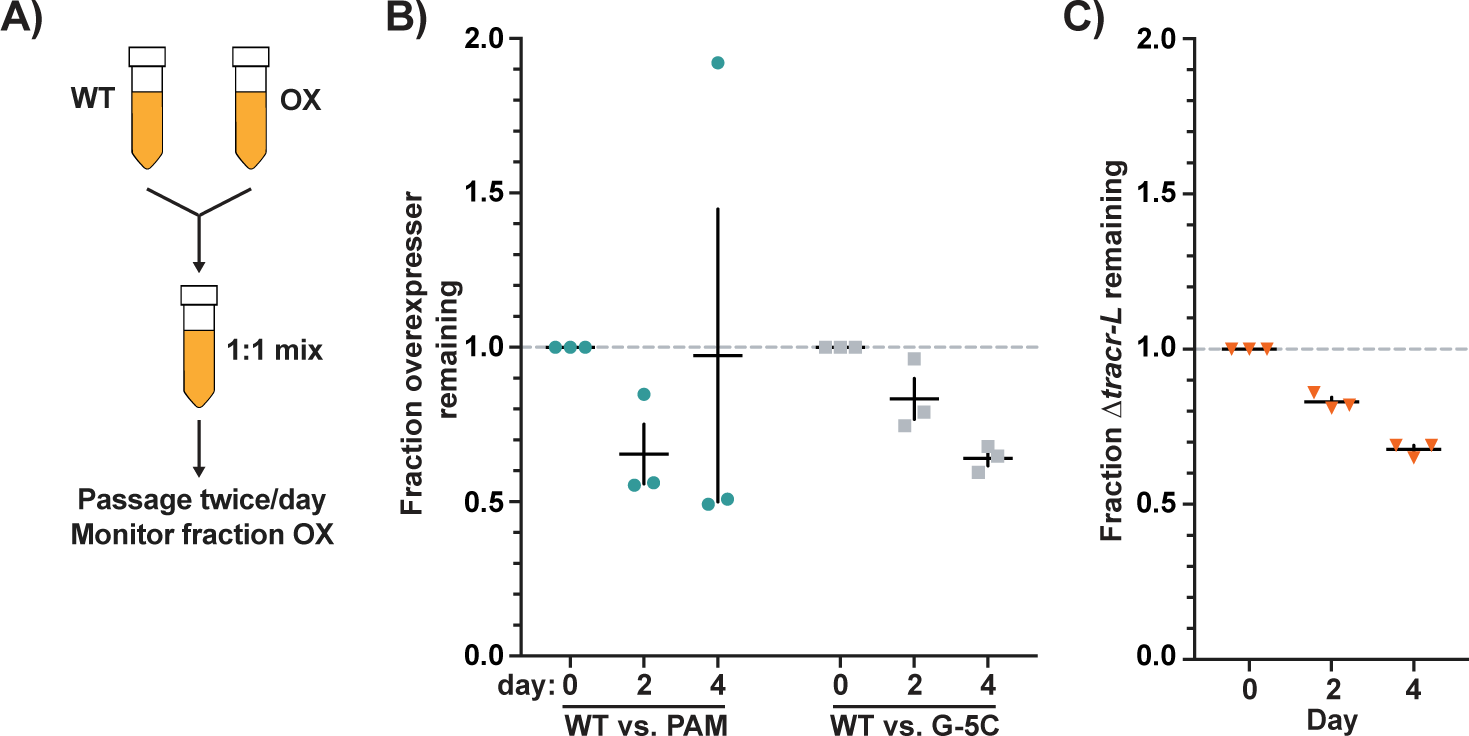
Overexpression of the *cas* operon decreases fitness. **A)** Schematic showing experimental setup for competition experiments. Equal amounts of overnight cultures were mixed and passaged every morning and evening for 4 days (8 total passages). The fraction of each strain in the coculture was quantified every two days. **B)** Competitions between WT SF370 and CRISPR-Cas PAM mutant or *tracr-L* target site mutant (G-5C) overexpressers. The fraction of overexpresser remaining relative to day 0 (dashed line) is shown. **C)** Competition between WT and Δ*tracr-L* cells. Errors bars are standard error. n=3.

We wondered whether Cas9 binding in other regions of the genome could explain the fitness costs we observed when CRISPR-Cas is overexpressed. This phenomenon has been observed in enterobacteria when spacers are introduced with short matches to essential gene promoters, leading to transcriptional repression and cytotoxicity^42^. To investigate this, we performed Cas9 ChIP-Seq to discover additional Cas9 binding sites along with RNA-Seq to determine whether genes surrounding those binding sites were differentially regulated in WT, Δ*tracr-L*, and ΔCRISPR-Cas cells. We found Cas9 bound to several regions containing partial matches to *tracr-L* or *crRNAs* (Fig. S4E, supplemental table 1). However, expression of genes surrounding these binding sites was not significantly altered by deletion of *tracr-L* or the full CRISPR-Cas system (supplemental table 2). Therefore, we do not think that specific transcriptional changes due to Cas9 binding explain the toxicity observed when CRISPR-Cas is overexpressed. Nonetheless, our results suggest that although CRISPR-Cas overexpression is beneficial during phage infection, it becomes detrimental to cells after the phage threat is cleared.

### Cas9:*tracr-L* binding is mutagenic

We were struck by the observation that, although mutations in either *tracr-L* RNA or its binding site generate a CRISPR-Cas overexpression phenotype, nearly all the overexpressers we sequenced had mutations in the binding site or PAM within P*_cas_* (Fig. 1C, S1B-C). Therefore, we wondered if Cas9:*tracr-L* binding elevates the rates of local mutagenesis. To test this, we constructed a strain of *S. aureus* (due to a lack of suitable inducible promoters in *S. pyogenes*) with the SF370 CRISPR-Cas system engineered into a genomic safe harbor^30^ with a *Δtracr-L* mutation and two plasmids carrying IPTG-inducible *tracr-L* cassettes. Thus, *tracr-L* was only expressed in this strain when IPTG was added. The CRISPR-Cas locus harbored a CRISPR array with a single spacer targeting phage φNM4γ4, and the dCas1 allele to prevent additional spacer acquisition. In the presence of IPTG, when *tracr-L* is present, this strain can only efficiently defend against φNM4γ4 at high MOIs if there is a mutation within P*_cas_* that disrupts *tracr-L* repression. Cells were passaged with and without IPTG to allow for mutations to occur over multiple generations (Fig. 5B). Then, all cultures were diluted into media with IPTG to induce *tracr-L* expression until Cas9 levels were equalized between the conditions. Cells were then infected with ɸNM4ɣ4 at MOI 100 in soft agar supplemented with IPTG (Fig. S5B). Strikingly, we found that inducing *tracr-L* during passaging resulted in a larger fraction of overexpressers in the surviving population (Fig. 5C, S5B-C). To rule out the possibility that growth in IPTG or *tracr-L* expression elevated global rates of mutagenesis, we also plated the same cultures on rifampin. *S. aureus* can evolve rifampin resistance through mutation of one of several residues within its target, RpoB, and rifampin is therefore frequently used to determine relative mutation rates^43–45^. The rate of rifampin resistance was not significantly different between the two populations (Fig. S5A), suggesting that the elevated rates of mutation we observed are specific to the Cas9:*tracr-L* binding site. Altogether, our results suggest that Cas9:*tracr-L* repressor binding is mutagenic and elevates the rate at which overexpressers are generated.

**Figure 5.**
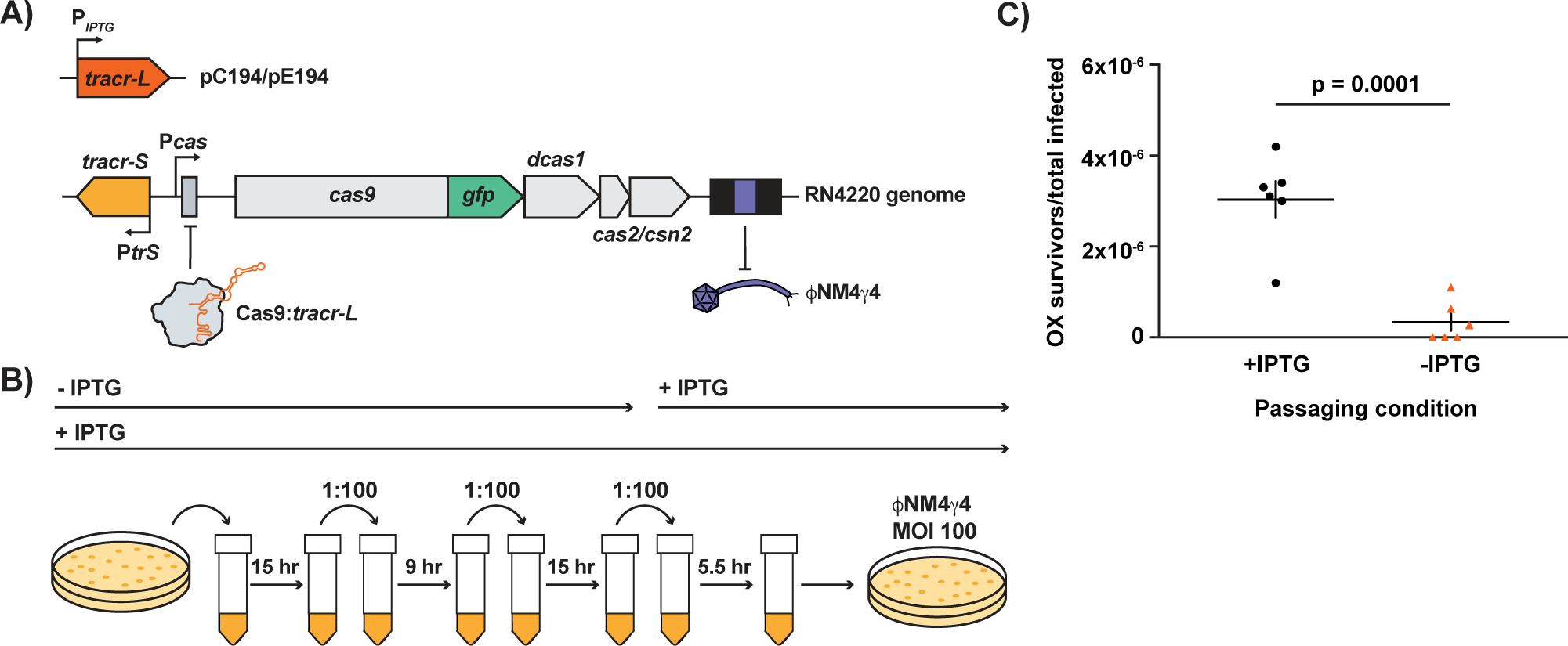
Cas9:*tracr-L* binding is mutagenic. **A)** Mutagenesis reporter strain. The *S. pyogenes* CRISPR-Cas system was integrated into the *S. aureus* RN4220 genome. *tracr-L* was expressed ectopically on two plasmids from an IPTG-inducible promoter. The CRISPR array contains one spacer targeting ɸNM4ɣ4. **B)** Experimental schematic. The reporter strain in (A) was passaged with (+ IPTG) and without (-IPTG) *tracr-L* induction. *tracr-L* expression was induced in all cultures before infecting with ɸNM4ɣ4 in soft agar. **C)** Colonies from phage infection in (B) were sequenced to determine if they had mutations in the *tracr-L* target site or PAM. The fraction of surviving OXs relative to the starting population is shown. Error bars are standard error. P-value determined using a two-tailed unpaired t-test. n=6.

## DISCUSSION

How bacteria regulate CRISPR-Cas expression to provide phage defense while avoiding autoimmunity remains an open question. There is a growing list transcription factors, both outside and within CRISPR-Cas loci, that regulate CRISPR-Cas expression in response to a wide variety of cues. For example, in the same *S. pyogenes* system described here, we recently found that phage-encoded anti-CRISPRs (Acrs) that inhibit Cas9 interference can also lead to transient inhibition of the Cas9:*tracr-L* repressive complex, producing a protective burst of CRISPR-Cas expression^9^. Here, we describe an alternative and complementary defense strategy, whereby a subpopulation of CRISPR-Cas overexpressing cells expands to defend the population against phage (Fig. 6). In contrast to transcriptional programs that sense and respond to phage, these hypervigilant cells express high levels of CRISPR-Cas components *before* the arrival of the phage and therefore provide highly efficient protection. At the population level, the enrichment of overexpressers functions as a rapid transcriptional response without requiring immediate regulatory activities in any individual cell.

**Figure 6.**
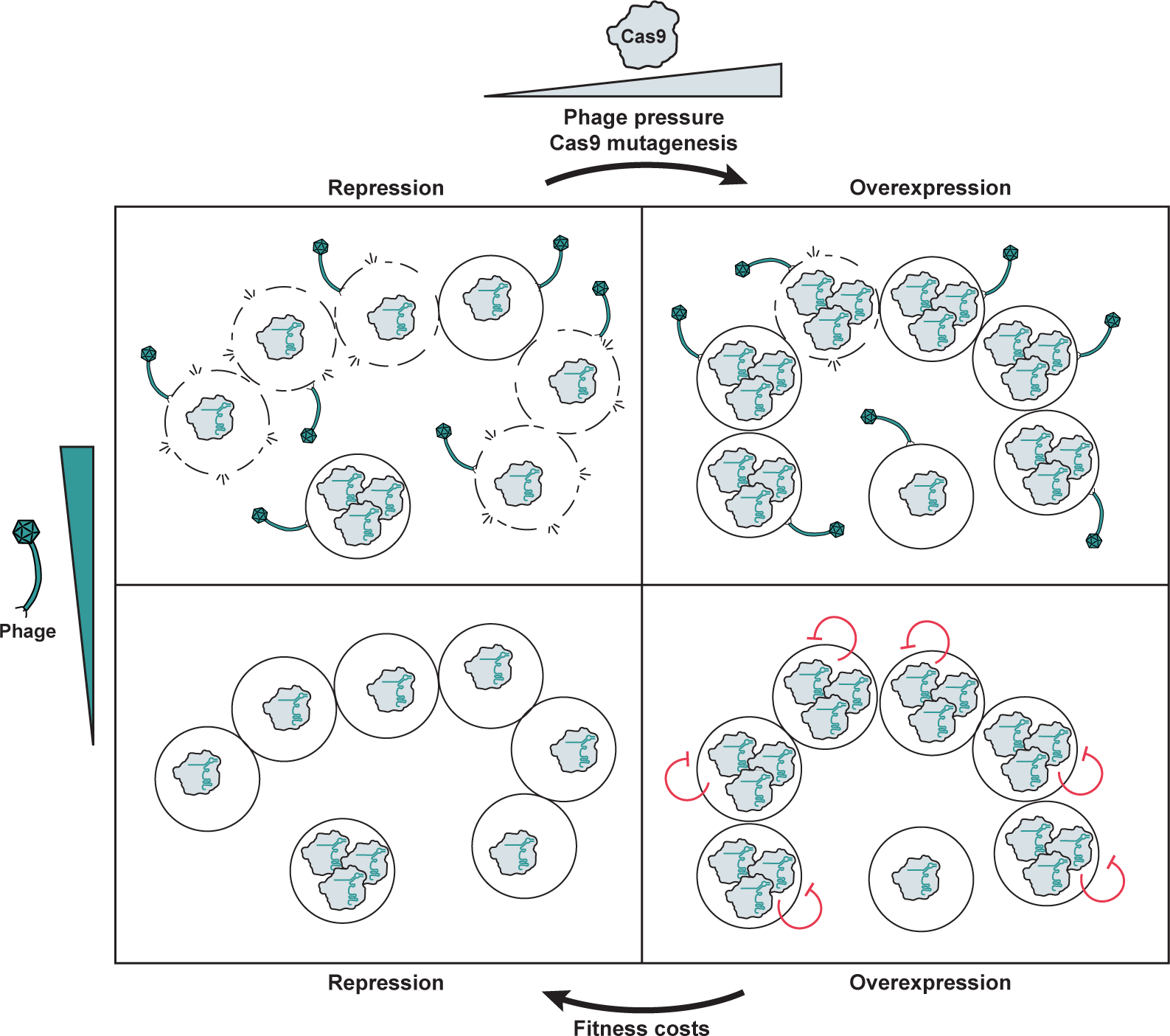
Competing pressures of phage infection and fitness costs lead to cycles of CRISPR-Cas repression and overexpression. A subpopulation of *S. pyogenes* overexpresses CRISPR-Cas and is enriched during phage infection when the majority of WT cells lyse. Cas9:*tracr-L* binding is mutagenic and increases the size of this overexpressing population. In the absence of phage, overexpressers are outcompeted due to fitness costs and CRISPR-Cas repression is restored. The cycle begins anew when the population is again threatened by phage.

We previously observed high levels of sequence variation in the *tracr-L* targeting sequence within the *Streptococci*^25^. Intriguingly, these mutations are accompanied by complementary changes in the P*_cas_* target site, maintaining base-pairing potential. Our results may help explain this covariation between *tracr-L* and P*_cas_* using a two-step model. First, cells harboring a mutation in P*_cas_* that leads to CRISPR-Cas overexpression are enriched during a phage infection. With enough phage pressure, these cells may take over the population. For example, we observed overexpressers go from ∼1/10 million to ∼1/10 of the overall population after an infection at MOI 1. After a similar infection with a second phage, overexpressers would be expected to take over the population. Second, after the phage threat has subsided, the fitness costs of overexpression favor cells that restore repression by either (i) mutating P*_cas_* back to the WT sequence or (ii) introducing a compensatory mutation in *tracr-L*, resulting in covariation.

For this strategy to be feasible, mutations leading to overexpression must occur frequently enough to maintain an overexpressing population. Here, we show that Cas9 binding leads to higher rates of mutations in the *tracr-L* target, thus elevating the rate at which overexpressers are produced. A similar phenomenon has been observed in yeast, where dCas9 binding increases mutation rates in and around the target site by 10-100 fold^46^. In biotechnology applications, this characteristic of Cas9 is undesirable. However, in its natural context, it may be an evolved feature rather than a bug. Cas9-induced mutagenesis may allow Cas9 to act as a switch which can both repress, and relieve repression of, immunity. Recently, many new examples of natural CRISPRi have been described, including mini arrays and *scaRNAs* that regulate genes throughout bacterial genomes^47–50^. A similar phenomenon could occur in those populations, and our data could more broadly explain how Cas9 repressors are dynamically regulated. In addition to Cas9 binding, some strains of *S. pyogenes* have a mutator phenotype due to genetic elements that eliminate DNA mismatch repair. In SF370, an integrated prophage-like element, SpyCIM1, disrupts *mutL*, leading to higher rates of mutation ^51,52^. This likely also elevates the rate at which mutations leading to CRISPR-Cas overexpression occur.

Our work highlights surprising constraints of the CRISPR array in WT *S. pyogenes* cells. Despite its modest size relative to other CRISPR arrays which can accumulate hundreds of spacers, only 3 of the first 4 spacers demonstrate potency against even the mild pressure of plasmid transformation. Curiously, the addition of a single new spacer severely dampened the efficiency of all other spacers, including those now occupying positions 2-4. These limitations suggest that CRISPR-Cas overexpressers may in fact play a significant, or even dominant role in phage defense. Overexpressers have an expanded effective array size of at least 9 spacers and can resurrect spacers that are dormant in WT cells. Additionally, overexpressers acquire spacers at a higher rate and frequently acquire multiple spacers, allowing them to sample more sequences and prevent the emergence of escaper phages.

Here we describe, to our knowledge, the first demonstration that CRISPR-Cas de-repression in a native host is associated with a fitness cost. This contextualizes the growing number of studies showing that CRISPR-Cas systems are natively repressed or silenced. ChIPSeq and RNASeq analysis suggest that the fitness cost is not due to a single Cas9 binding site that alters gene expression. However, we cannot rule out that Cas9 binding leads to subtle transcriptional changes below our limit of detection which nevertheless are detrimental over generations. Alternatively, transient Cas9 binding events at off-target sites could lead to DNA damage, either by Cas9 cleavage or collisions with the transcriptional or replication machinery. While the precise mechanisms of autoimmunity will require further study, our work emphasizes that CRISPR-Cas toxicity is not simply a consequence of overexpression in a heterologous host. Intriguingly, in *Streptococcus thermophilus*, there is a fitness cost associated with even basal levels of CRISPR-Cas expression that is relieved when *cas9* or *csn2* are knocked out, although it is unclear whether or not this system is natively transcriptionally regulated^53^.

Recently, connections have been drawn between newly discovered bacterial phage defense systems and vertebrate innate immunity, with many cell-autonomous immune effectors demonstrating similar functions and even shared evolutionary history^54^. Although prokaryotic and vertebrate adaptive immune effectors are not evolutionarily related, they share mechanistic and functional similarities. In both cases, memories are stored genetically, as spacers in the CRISPR array or as coding sequences for specific antibodies in VDJ loci, respectively. In addition, the memory formation process depends upon recombinases derived from transposons (Cas1 in prokaryotes, and Rag1-Rag2 in vertebrates)^55,56^. One fundamental difference is that CRISPR-Cas memory formation is one of few examples of *bona fide* Lamarckian evolution^57^, whereby the genome is modified in response to an environmental change (phage infection). However, here we describe a bacterial adaptive immune strategy conforming to Darwinian evolution; phage selection acts on preexisting variation, leading to expansion of the CRISPR-Cas overexpressing population. This is not unlike the process of B-lymphocyte clonal selection and expansion in response to vertebrate pathogens, yet another striking similarity between immune strategies in organisms from different domains of life.

## SUPPLEMENTAL FIGURES

**Figure S1.**
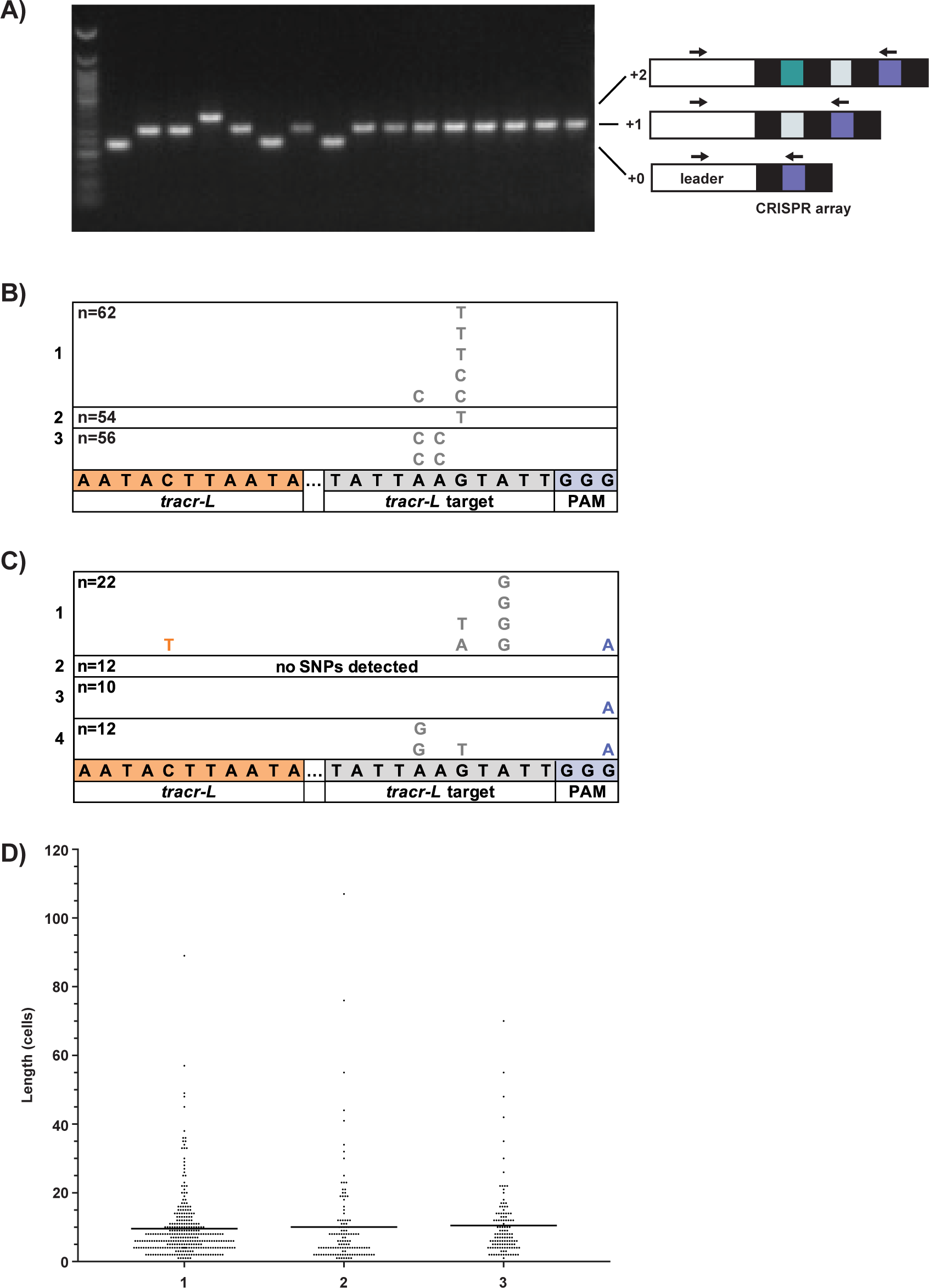
A subpopulation of CRISPR-Cas overexpressers expands during phage infection. **A)** Primers that bind to the leader sequence and spacer 1 were used to check for spacer acquisition in survivors of A1 infection (Fig. 1B). The size of the PCR product (representative agarose gel shown) was used to determine the number of spacers acquired by each survivor. **B)** Mutations observed in the *tracr-L* target sequence as summarized in Fig 1C, separated by replicate. ‘n’ refers to the total number of survivors screened per replicate for spacer acquisition and SNPs. **C)** Mutations in *tracr-L* regulatory sequences from surviving dCas1 colonies, separated by replicate. ‘n’ refers to the total number of survivors sequenced per replicate. **D)** Number of SF370 cells in individual chains plotted for three biological replicates. The average chain length for all replicates is 10 cells. n = 130 – 300 chains per replicate.

**Figure S2.**
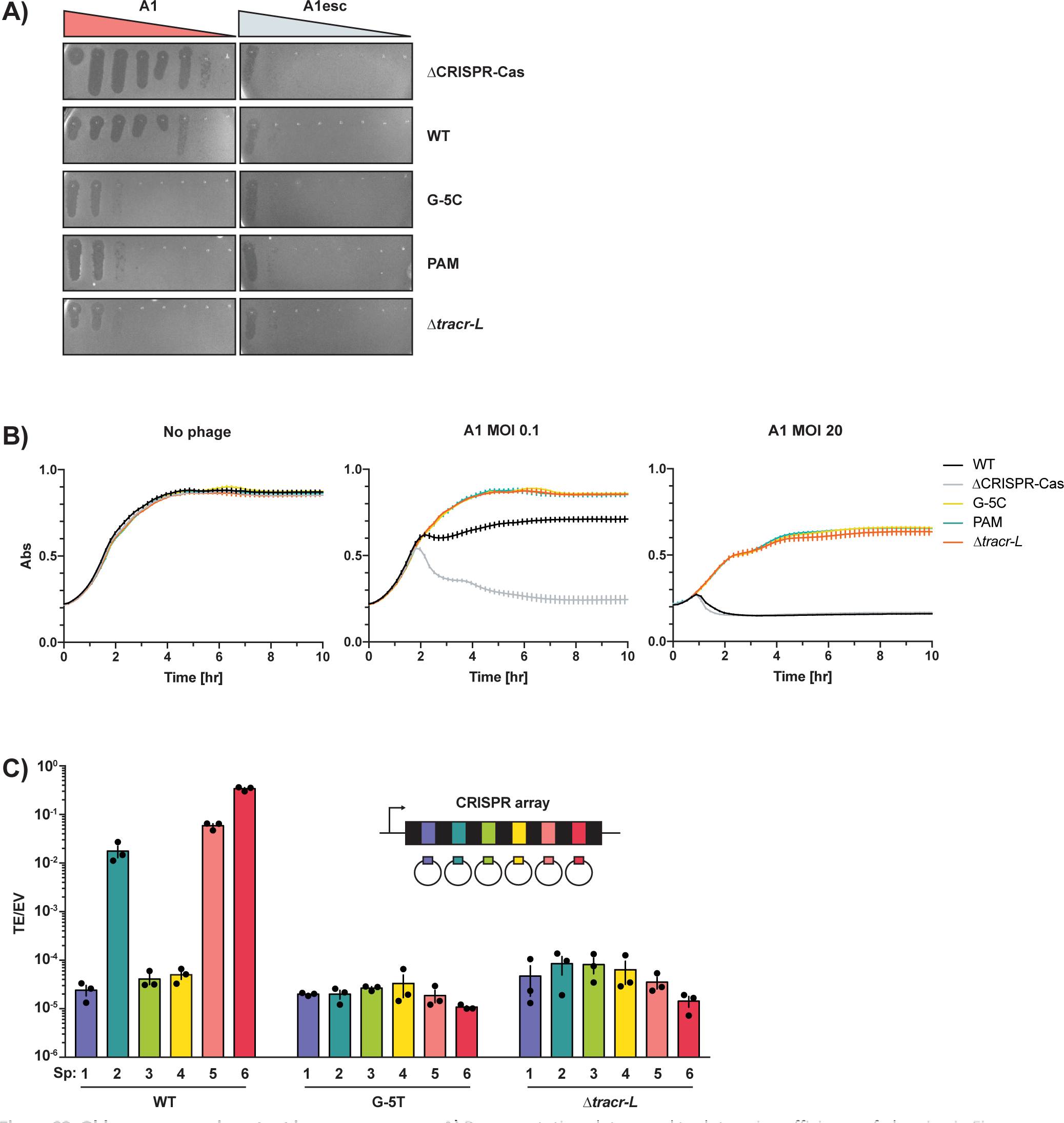
Old spacers remain potent in overexpressers. **A)** Representative plates used to determine efficiency of plaquing in Fig. 2A. **B)** Liquid infections of cells with mutations in *tracr-L* regulatory sequences at MOI 0, 0.1, and 20. **C)** WT and CRISPR-Cas overexpressing cells were transformed with plasmids containing protospacers targeted by each of the 6 spacers in the native CRISPR array. Transformation efficiency was calculated relative to an empty vector. Error bars are standard error. n=3.

**Figure S3.**
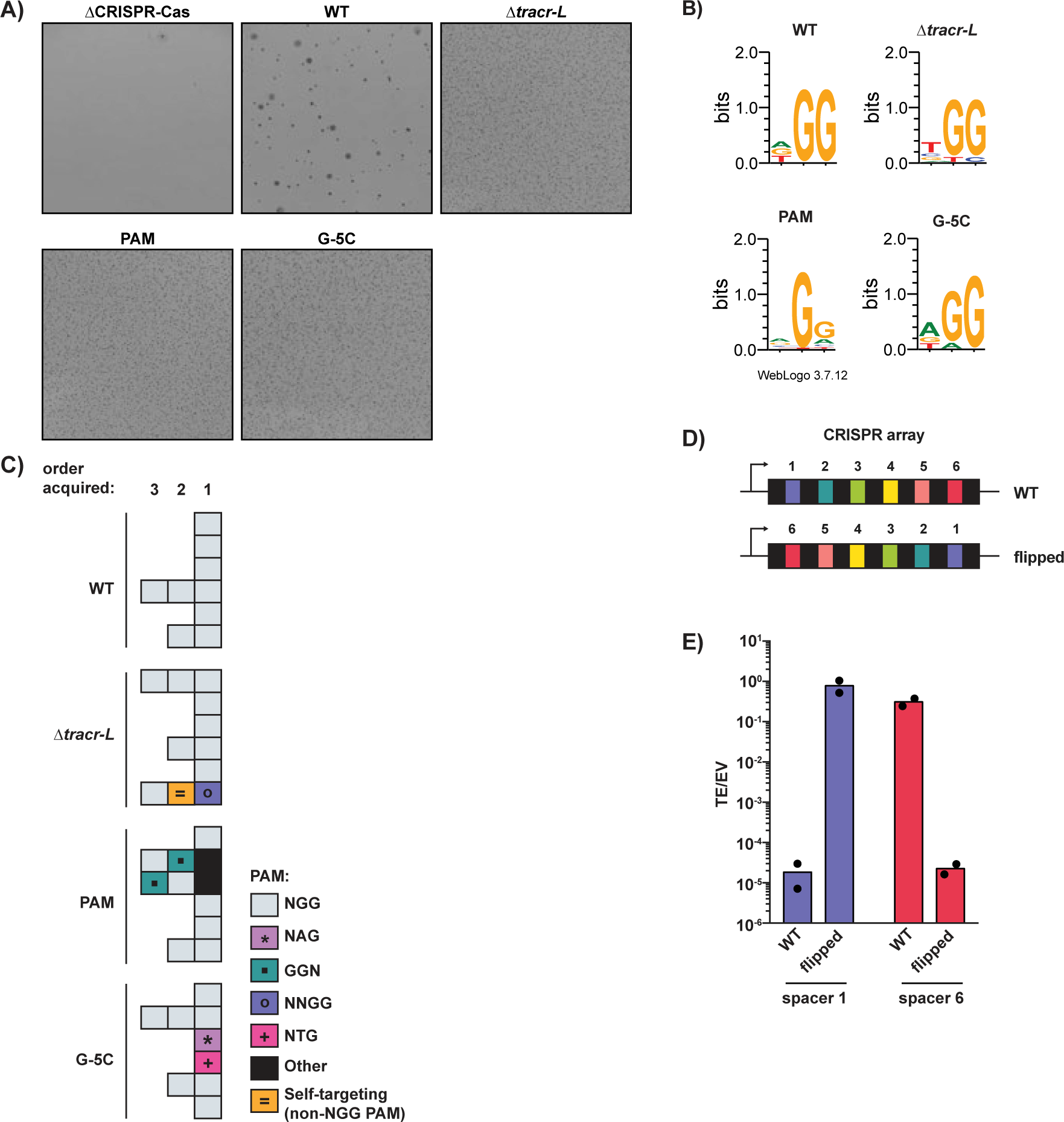
CRISPR-Cas overexpressers generate greater functional spacer diversity. **A)** Representative views of colonies on soft agar plates from immunity assay in Fig 3A. **B)** Spacers acquired in six surviving colonies of each strain from (A) were sequenced, and their targets were identified by alignment to the phage A1 or SF370 genome. Sequence logos of all sequenced protospacer PAMs are shown. **C)** CRISPR array architecture in colonies described in (B). Each row represents the new spacers acquired in one colony, where the leftmost box is the newest, leader-proximal spacer. The nucleotide identity of the protospacer PAM is represented by the color of and symbol within each box. **D)** Order of spacers in WT SF370 and a strain in which the spacers were flipped (Note: the 5’-3’ orientation of each spacer sequence is maintained). **E)** Transformation efficiency of plasmids targeted by the first or last spacer in the SF370 array transformed into WT cells, and a strain with the order of spacers flipped.

**Figure S4.**
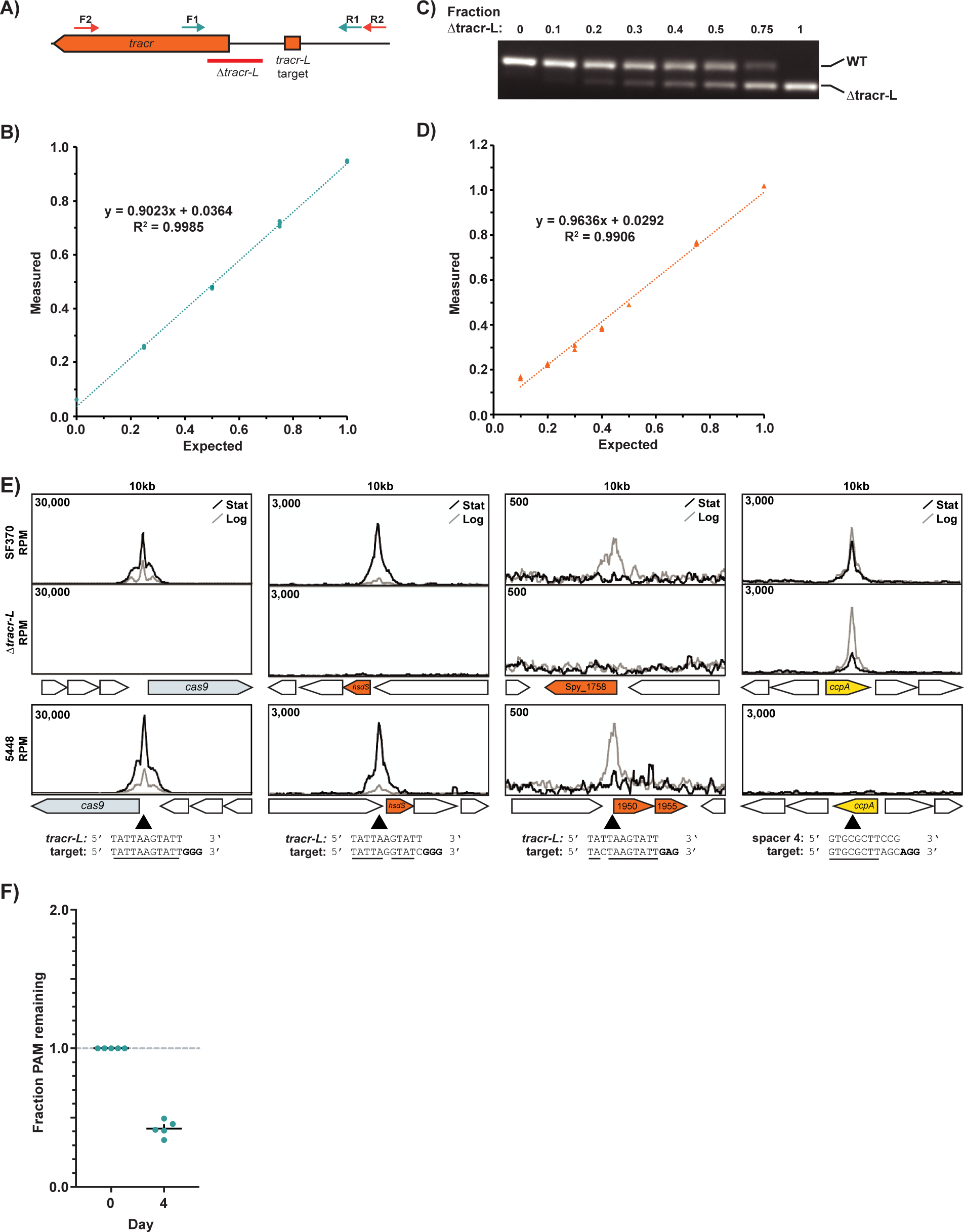
Overexpression of the *cas* operon decreases fitness. **A)** Binding sites of primers used for amplicon sequencing (F1/R1) and semi-quantitative PCR (F2/R2) to quantify fractions of overexpressers in competition assays. **B)** Validation of amplicon sequencing strategy to quantify WT vs. PAM/target site mutant competitions. The region containing the *tracr-L* target site and PAM was amplified and sequenced with Next Generation Sequencing. The fraction of reads corresponding to each genotype was quantified. n = 2 for all fractions except 0 and 1, where n=1. **C)** Validation of semi-quantitative PCR to quantify WT vs. Δ*tracr-L* competitions. PCR across the *tracr* locus was performed using mixtures of known fractions of WT and Δ*tracr-L* cells as template, resulting in a longer product from WT cells and a band 52bp shorter from Δ*tracr-L* cells, and run on agarose gels. **D)** The fraction of Δ*tracr-L* cells from (C) was quantified by dividing the normalized Δ*tracr-L* band intensity by the combined intensity of WT and Δ*tracr-L* bands. n = 2 for all fractions except 1, where n=1. **E)** Cas9 ChIP data from stationary (stat) and logarithmic (log) phase cultures are shown at P*_cas_* and partial matches to *tracr-L* and *crRNA*s in SF370, an SF370 Δ*tracr-L* strain, and the M1 GAS strain 5448. Windows are centered around the target site (black arrow), and the sequence of the target and its guide RNA are shown below. Matching bases are underlined, and PAM sequences are bolded. In strain 5448, the sequence of *tracr-L* and all target sites shown are perfectly conserved, but spacer 4 is not present in the CRISPR array. n = 1. **F)** Results of competitions between WT and PAM mutant cells. The fraction of PAM mutants remaining on day 4 is shown normalized to the starting fraction. Error bars are standard error. n=5.

**Figure S5.**
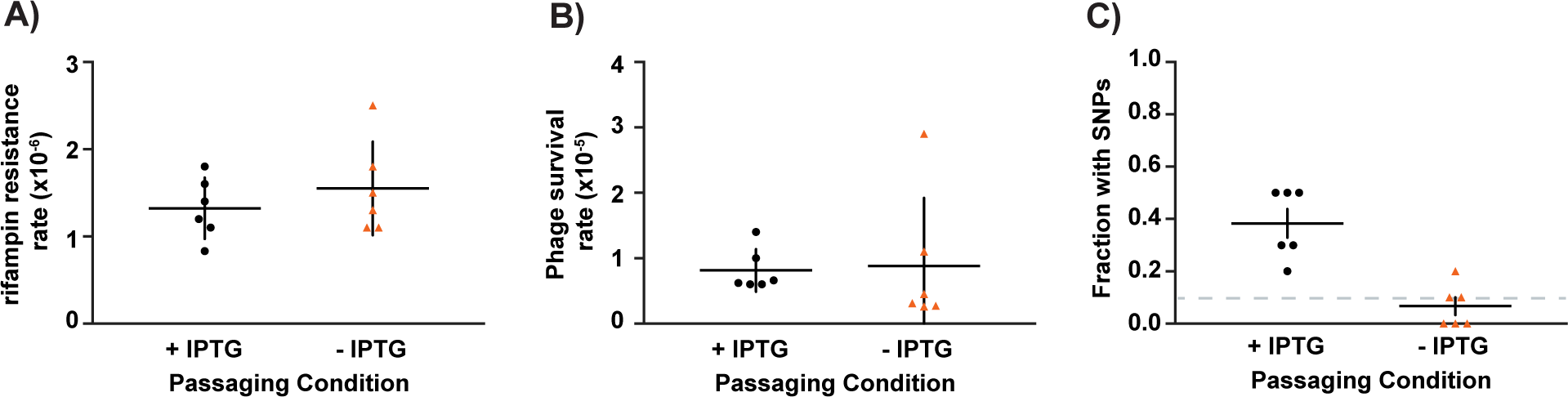
Cas9:*tracr-L* binding is mutagenic. **A)** Rifampin resistance rates of populations passaged with (+ IPTG) and without (-IPTG) *tracr-L* expressed. **B)** Survival rate of cells passaged with and without *tracr-L* expression when infected with ɸNM4ɣ4 at MOI 100 in soft agar. **C)** The fraction of sequenced colonies with mutations leading to CRISPR-Cas overexpression from plates described in (B). Dotted line indicates limit of detection. Error bars are standard error. n=6.

## Notes

### Competing Interest Statement

The authors have declared no competing interest.

